# Systemically administered MSCs given 24hrs after osteotomy do not affect bone formation in rat distraction osteogenesis

**DOI:** 10.1101/293514

**Authors:** Jonathan Guevara, Zacharie Toth, Daniel Kim, John Peters, Adrian Marley-Weaver, J Tracy Watson, Sarah McBride-Gagyi

**Author notes:** **Corresponding Author:** (SM).

## Abstract

Distraction osteogenesis is a unique postnatal bone formation employed by orthopaedic surgeons to treat many conditions, however, the overall time to external frame removal can be extensive. Any strategies that accelerate healing would improve patient care. Distraction osteogenesis research in the past decade has shown that direct stem cell implantation enhances new bone formation. Systemic implantation would be more clinically desirable. Systemically delivered stem cells have been shown to home to a mandibular distraction site; however, effects on bone formation have not been studied. Ten-week-old, male Sprague-Dawley rats underwent surgery to implant an external fixator-distractor and an osteotomy was performed. Twenty-four hours postoperatively, each rat received tail vein injections of either saline or 10^6 fluorescently labeled primary mesenchymal stem cells. Animals in the validation groups were euthanized two days after surgery and the femora processed for histology. Animals in the experimental groups were given five days of latency, then the femur was lengthened once daily for five days (0.75mm/day, 3.75mm total). Following four weeks of consolidation, the animals were euthanized and the femora were evaluated by microCT and histology to quantify new bone formation. Labeled stem cells were found at the distraction site in validation animals. However, there were no differences in any bone or soft tissue outcomes. Systemic stem cell administration 24 hours after surgery does not improve DO outcomes. It is possible that the animal model was not challenging enough to discriminate any augmentation provided by stem cells.

## Introduction

Distraction osteogenesis (DO) is a form of post-natal bone formation that is frequently employed by orthopaedic surgeons to treat a variety of conditions. This unique and powerful technique has been in use for several decades, and was propagated by Ilizarov, borrowing from decades of fracture care and limb lengthening procedures.[1–3] Currently, DO is clinically applicable in situations of limb salvage, segmental bone defects, angular deformities of childhood, malunions/nonunions in the trauma population and limb length discrepancies.[1,3–6] There are four phases[3,5]: surgery, latency, distraction and consolidation. During the surgical phase, an external or internal fixation device is applied to the bone that allows distraction and provides mechanical stability during the subsequent phases. A corticotomy is then performed in an area of healthy bone. The latency period then follows prior to the initiation of distraction. This latency period allows time to initiate a bone repair response (typically 5-7 days). During the distraction phase the corticotomy is slowly distracted at a maximal rate of 1 millimeter per day. by elongating the external fixator or internal device. Once the desired new length is reached, distraction is stopped and the consolidation phase begins. During consolidation, the distracted osteogenic tissue mineralizes and remodels until the external fixator can be removed. Typically this phase lasts from 1 to 2 months per centimeter of lengthening. As widely used as this technique has become, the protracted time periods for the distraction and consolidation phases limit its clinical utility.[1,3,7–10] Strategies to speed up either of these two phases would decrease treatment time and improve patient quality of life.

A body of pre-clinical research over the past 20 years indicates that addition of cells enhances the amount and quality of new bone generated by DO.[11–25] These studies have used stem cells or osteogenically differentiated cells with success. This has been shown in both mandibular[12,13,16–19,25,26] and long bone distraction[11,14,15,20–24] and could theoretically decrease the consolidation time. Recent studies even indicate that trophic factors released by stem cells allow faster rates of distraction than previously attainable.[20,27] Interestingly, stem cell treatment positively augments bone generation regardless of which phase it is administered although few studies have tested that hypothesis. However, all studies to date applied the cells or factors directly to the osteotomy or newly distracted tissue.

In the clinical setting, directly applying stem cells may not desirable for several reasons. First, injection to the osteotomy site could disrupt the newly forming bone especially if the patient has comorbidities such as smoking not present in animal models. Second, to avoid injection errors, stem cells would need to be administered by a surgeon with some level of anesthesia. Consequently, unless injection is done during the initial surgical osteotomy or before release from the hospital, success relies on the patient to return at the appropriate time and undergo an additional anesthesia event. Lastly, there is the issue of cell source. Application at the time of surgery entails either donor stem cells or previously harvested autologous stem cells. Using donor stem cells requires some level of immunosuppression, while preemptive autologous harvest from long bone sources necessitates an additional anesthesia event. Harvest of autologous cells at the time of surgery followed by systemic administration via intravenous route would afford the same theoretical advantage as direct implantation; with minimal additional risk to the patient or the distraction site. Furthermore, it may be advantageous to introduce stem cells the day after surgery at a time point when natural chemokine gradients are forming and the inflammatory cascade is at its peak.[8,9,26] During this time, migration and recruitment of regenerative cells is very high.

Mesenchymal stem cells (MSCs) have been found to migrate to fracture or drill hole sites demonstrating a positive osteogenic effect.[28–39] Homing and implantation capabilities are enhanced when cells are applied 24 hrs after injury as opposed to later time points.[35] Studies have shown that animals that receive systemic MSCs have increased bone volume,[30,32,37] and accelerated callus mineralization.[30,33,34,40] Combined this leads to improved regenerate formation and greater mechanical strength.[30,37,40] Systemically-administered MSC behavior during DO has not been extensively studied. To our knowledge there is only one study that investigates this topic. Cao *et al.*[41] proved that systemically applied MSCs will home to the newly distracted tissue in a rat mandible. Furthermore, up- or down-regulation of the CXR-L and SDF-1 pathway, which regulates stem cell homing, will increase or decrease the number of exogenous MSCs that home to the distraction site, respectively.[41] However, this study did not examine bone formation. It is unknown if systemically-administered MSCs will enhance bone regeneration during DO similar to direct implantation.

Given the positive effects of directly implanted MSCs on DO and the MSC’s ability to home to the distraction site, we hypothesize that systemic administration of MSCs will improve bone formation in DO. The purpose of this study was to verify that systemically injected MSCs will home to the site of a femoral distraction in a rat model and to investigate if systemic application would positively augment bone formation. Since human bone repair is similar to that of rodents, a rat model of femoral distraction osteogenesis was employed to test the hypothesis. Experimental rats were given a tail vein injection of stem cells 24 hrs after surgery. After 5 days of distraction and 4 weeks of consolidation, the limbs were harvested to quantify the newly formed bone in the distraction site via microCT and the soft tissue types via histology.

## Materials and Methods

### Animals

All experiments were carried out with the approval of our institution’s IACUC (Protocol Number: #2427). Rats were kept in standard husbandry conditions with light/dark cycles and fed *ad libitum*. Eight-week-old male Sprague-Dawley rats (SAS SD, Charles River, Wilmington, MA, USA) were used for cell harvesting. Cohort ten-week-old male Sprague-Dawley rats were used for validation and distraction experiments. Prior to surgery animals were group housed. After surgery they were single housed to prevent damage to sutures or installed hardware. Euthanasia was carried out via carbon dioxide asphyxiation using AVMA recommended guidelines (graduate replacement at 10-30% chamber volume/min).

### Stem Cell Isolation and Labeling

MSCs were harvested from the femora and tibiae of non-operative cohort rats for systemic transplantation. Briefly, rats were euthanized and the femora and tibias dissected out. After removing the surrounding soft tissues and articulating ends, the marrow was agitated with an 18 gauge needle and removed by centrifugation using nested micro-centrifuge tubes. This collected marrow was diluted into culture media (Minimal Essential Media, MEM supplemented with 20% Fetal Bovine Serum, 1% penicillin-streptomycin, and 1% L-glutamine) and the cells homogenized by repeated, gentle flushing with an 18 gauge needle and syringe. The suspension was filtered through a 70 μm cell strainer followed by a repeat cycle of centrifugation and homogenization.

Cells were then counted, plated into T-75 cell culture flask, and cultured in standard conditions (37°C and 5% CO_2_). Due to the large number of cells placed in a single culture flask, after 36 hours the supernatant was removed into a new T-75 flask to capture any remaining plastic adherent cells that could not find adequate space in the original flask. The media in the original flask was replaced. The culture media in both flasks was replaced every 24 hours. Seven to 9 days of culture were needed to obtain sufficient cells to inject 3-4 rats. Shortly before time of injection the cells were lifted from the culture flask surface by incubation with trypsin (5%, 37C, 5min), pelleted, and resuspended in suspended in sterile saline at a concentration of 2x10^7^ cells/mL. An equal volume of 2mM CSFE (5(6)-Carboxyfluorescein N-hydroxysuccinimidyl ester, Sigma) in 1X PBS was added and incubated for 10 minutes to fluorescently label the cells. Cells were then rinsed in serum rich PBS to removed excess CSFE, counted, and the concentration adjusted such that 10^6 cells could be injected in 400 to 500uL of sterile saline.

### Surgery & Stem Cell Administration

A surgical model of rat femoral DO was used. First, rats were anesthetized with isoflurane gas (3.5% for induction, 2.0% for maintenance). The skin overlying the right femur was shaved and disinfected with chlorhexidine prior to incision. An incision from the greater trochanter to the lateral femoral condyle was made to expose the lateral surface of the femur. Soft tissue dissection was carried out exploiting the interval between the vastus lateralis and the biceps femoris, and the anterolateral surface of the femur was exposed. Four 1.2mm pilot holes were drilled through both cortices with a power drill (8V Variable Speed Lithium Ion Drill, Ryobi, Anderson, SC) using the custom external fixator as a guide. Then fully-threaded Kirschner-wires (K-wires, 1.25 inch long, 0.062 inch diameter, Microaire, Charlottesville, VA) were implanted, and the external fixator was secured 10mm from the bone surface.

With the external fixator in place, a 0.60mm osteotomy was manually created at the bone mid-shaft using a 41 teeth per inch scroll saw blade (Isomax, Easypower Corporation, Chicago, IL), and the limb was shortened to appose the bone ends. The overlying skin was closed with nylon 5-0 suture (McKesson, Richmond, VA). Prophylactic antibiotics (Baytril, enrofloxacin, 10mg/kg BW, intramuscular route) were given the day before, the day of, and within 3 days following surgery. To alleviate pain a single injection of Buprinex (Buprenorphine SR-lab, 1mg/kg BW, subcutaneous route) was given just prior to surgery. This sustained release formula provides effective pain relief at least 72hrs. Twenty-four hours after surgery, the either saline (500uL) or 10^6 MSCs was administered systemically via tail vein injection based on previous dosage studies.[32] Animals were randomly assigned to each group.

### Stem Cell Homing Validation

Six rats (n=2 saline, n=4 MSCs) were used to verify stem cell homing. Twenty-eight hours following systemic injection, the rats were euthanized (post-operative day 2). The operative limb and surrounding soft tissues were harvested and placed into 10% neutral buffered formalin for 72 hours. Following fixation the distractor device was removed, and the tissues were decalcified with formic acid (immunocal, Statlab, McKinney, TX) for 4 days. The limbs were cut longitudinally along the sagittal plane, infiltrated with 30% sucrose, and embedded in OCT media for cryosectioning. Sections were counter stained with DAPI and the distraction site imaged in blue and green channels to visualize the cell nuclei and labeled cells, respectively (Leica DM4100B with DFC340FX Camera, Leica Systems). Images were qualitatively accessed for the presence (or absence) of cells in the MSC treated samples.

### Bone Augmentation

Following surgical application of external fixator and a latency period of 5 days, the bone was lengthened by 3.75mm (0.75mm/day for 5 days) (n=8 saline, n=10 MSC) based on previous studies.[11,14,21,22] A single injection of Buprenex was given by subcutaneous route for pain relief on the first and fourth day of distraction. Maintenance of alignment and callus maturation were monitored via biweekly radiographs starting on the final day of distraction. Experimental animals were euthanized at 4 weeks of consolidation based on available literature and our pilot studies.[11,15,23] Distracted limbs were then harvested, fixed and processed for microCT and histology.

### MicroCT

MicroCT was used to determine the amount, structure and quality of the newly formed tissue. The distracted limb was scanned to encompass the entire distracted area and some adjacent bone (Fig 1A, white box) (MicroCT 35, ScanCo Medical; X-ray tube potential 70 kVp, integration time 300 ms, X-ray intensity 145 µA, isotropic voxel size 10 um, frame averaging 1, projections 1000, high resolution scan). The volume of interest for each sample was defined as the most proximal to the most distal slice that the endochondral cortex was disrupted excluding the preexisting cortical bone. Two different analyses were performed on each scan. In the first analysis, contours were drawn to encompass only newly formed bone (Fig 1B, black lines). This was done to avoid unbridged defect areas diluting outcomes dependent on total space (e.g. trabecular number per unit volume). This method was used to measure bone volume (BV), trabecular thickness (Tb.Th), number (Tb.N), spacing (Tb.Sp), connection density (Tb.Con), bone mineral density (BMD), and tissue mineral density (TMD). In the second method, contours included any area where bone was expected (i.e. both bone and unbridged space) (Fig 1C, black lines). The new tissue’s total volume (TV) and BV/TV were calculated from this method.

**Fig 1.**
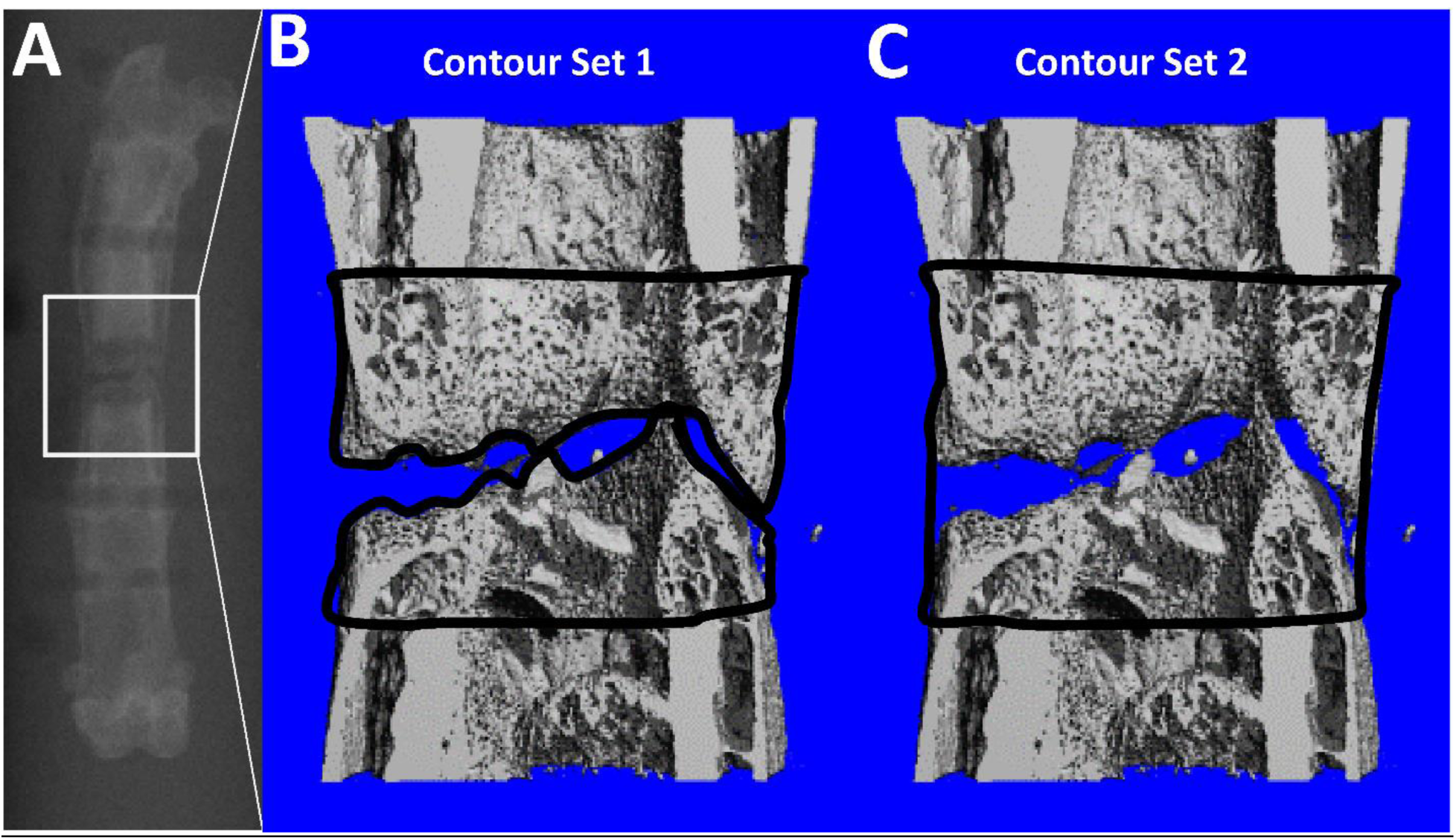
MicroCT Analysis Example. (A) The area encompassing the distracted area and some of the original bone ends was scanned at a 10um resolution (white box). **(B)** The first contour set (black lines) included only newly formed bone so that bone architecture outcome parameters dependent on total volume (e.g. trabecular number per unit volume) were not artificially diluted by the inclusion of unbridged space. **(C)** The second contour set (black line) included the entire area that bone would be expected if fully bridged in order to represent the completeness of healing.

### Histology

After being scanned by microCT, the samples were processed for histology (n=6 Saline, n=9 MSC). Briefly they were decalcified in formic acid for 4 days, longitudinally cut along the sagittal plane, embedded in paraffin, and sectioned. Sections were stained with hematoxylin, picrosirius red, and Alcian blue to differentiate cell nuclei, bone and fibrous tissue, and cartilage respectively and imaged (Leica DM4100B with DFC295 Camera, Leica Systems). The images were analyzed with a custom MATLAB program (Mathworks, Natick, MA) to quantify the amount of bone, cartilage, fibrous tissue, and marrow in the defect space.

### Statistics

Data shown is mean plus or minus standard deviation. Each outcome was compared between groups using unpaired t-tests. A p-value of <0.05 was considered significant. (Statview, v5.0, SAS Institute, Inc.)

## Results

### Validation of MSC Homing

Fluorescent microscopy was used to visualize the CFSE labeled cells in cryosections 1 day after injection. Saline injected controls had no green signal in the osteotomy. Labeled cells were found in the osteotomy and surrounding tissues of all MSC injected animals (Fig 2). There appeared to be a gradient with the highest cell concentration at the osteotomy and surrounding the K-wires, but this was not quantified.

**Fig 2.**
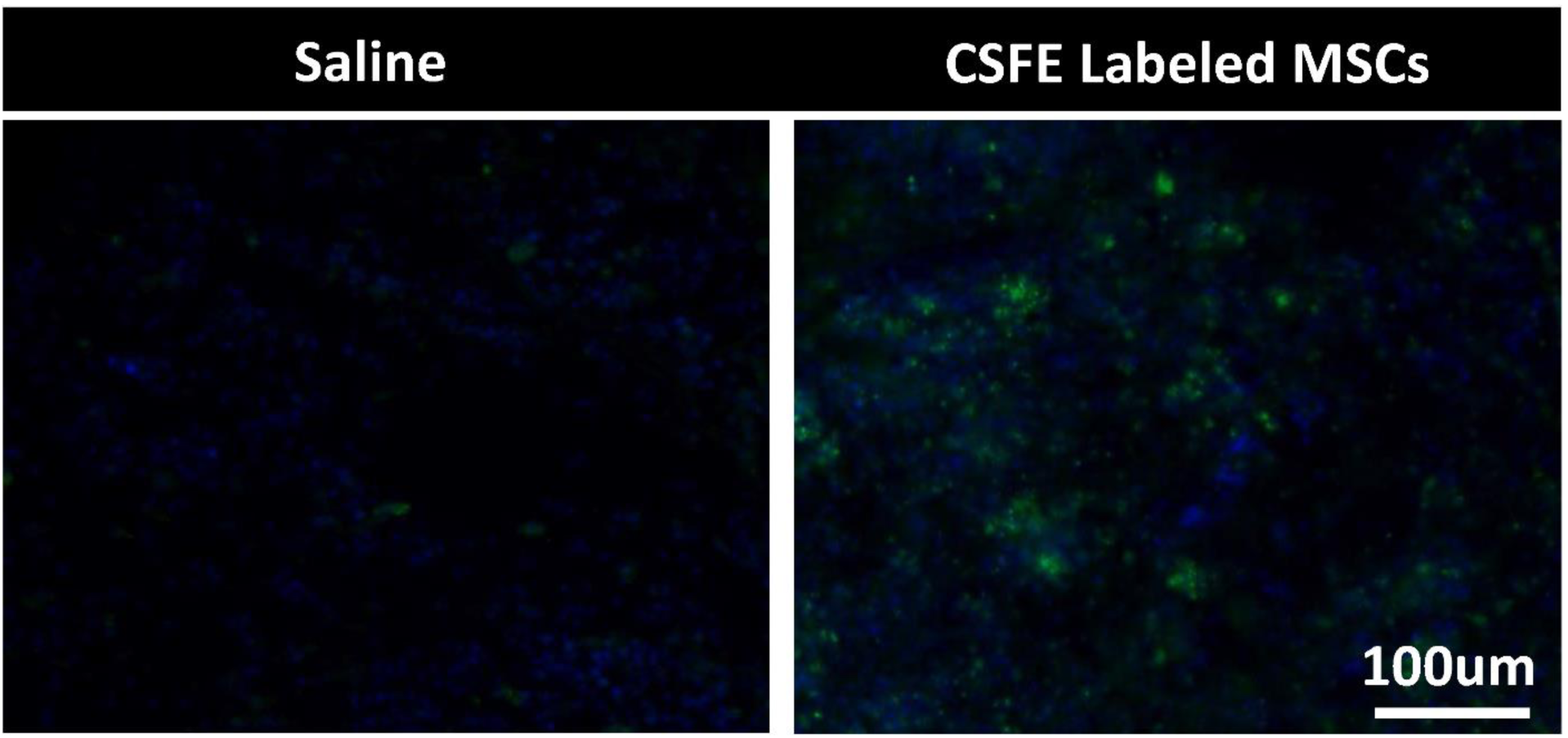
Stem Cell Homing. Images taken from the center of the osteotomy 28 hours after tail vein injection (post-op day 2). Saline injected rats had little to no green signal in the defect while MSC injected rats had noticeable green signal demonstrating that labeled MSCs had migrated from the circulatory system to the injury site.

### Quantity and Quality of bone formation

MicroCT was used to access the volume, density, and structure of the newly formed distracted bone after 4 weeks of consolidation (Table 1, Fig 3). Histology was used to access soft tissues at the same time point (Table 1, Fig 4). All samples had new bone formation which was trabecular in structure; however only one in each group was bridged across the entire gap. There were no differences between groups for any microCT or histology outcome.

**Table 1.** MicroCT and Histology Results

|  | Outcome | Unit | Group |  |  |  |
| --- | --- | --- | --- | --- | --- | --- |
|  |  |  | Saline |  | MSC |  |
|  |  |  | Mean | Std | Mean | Std |
| MicroCT | TV | mm<sup>3</sup> | 60.6 | 11.4 | 62.9 | 20.1 |
|  | Bone Volume BV | mm<sup>3</sup> | 20.2 | 5.6 | 20.0 | 8.6 |
|  | BV/TV | - | 0.332 | 0.062 | 0.316 | 0.083 |
|  | Tb.Th | mm | 0.088 | 0.011 | 0.082 | 0.014 |
|  | Tb.N | 1/mm | 4.62 | 1.52 | 4.45 | 1.10 |
|  | Tb.Con | 1/mm<sup>3</sup> | 385.0 | 123.6 | 454.1 | 124.3 |
|  | Tb.Sp | mm | 0.316 | 0.171 | 0.300 | 0.092 |
|  | BMD | mgHA/cm<sup>3</sup> | 435.0 | 61.0 | 414.7 | 91.1 |
|  | TMD | mgHA/cm<sup>3</sup> | 957.4 | 23.8 | 958.4 | 22.4 |
| Histology | Percent Marrow | % | 48.7 | 10.6 | 56.8 | 5.9 |
|  | Percent Fibrous | % | 5.0 | 5.2 | 5.9 | 4.7 |
|  | Percent Cartilage | % | 4.6 | 3.8 | 6.3 | 6.8 |
|  | Percent Bone | % | 47.4 | 9.5 | 41.1 | 5.5 |

**Fig 3.**
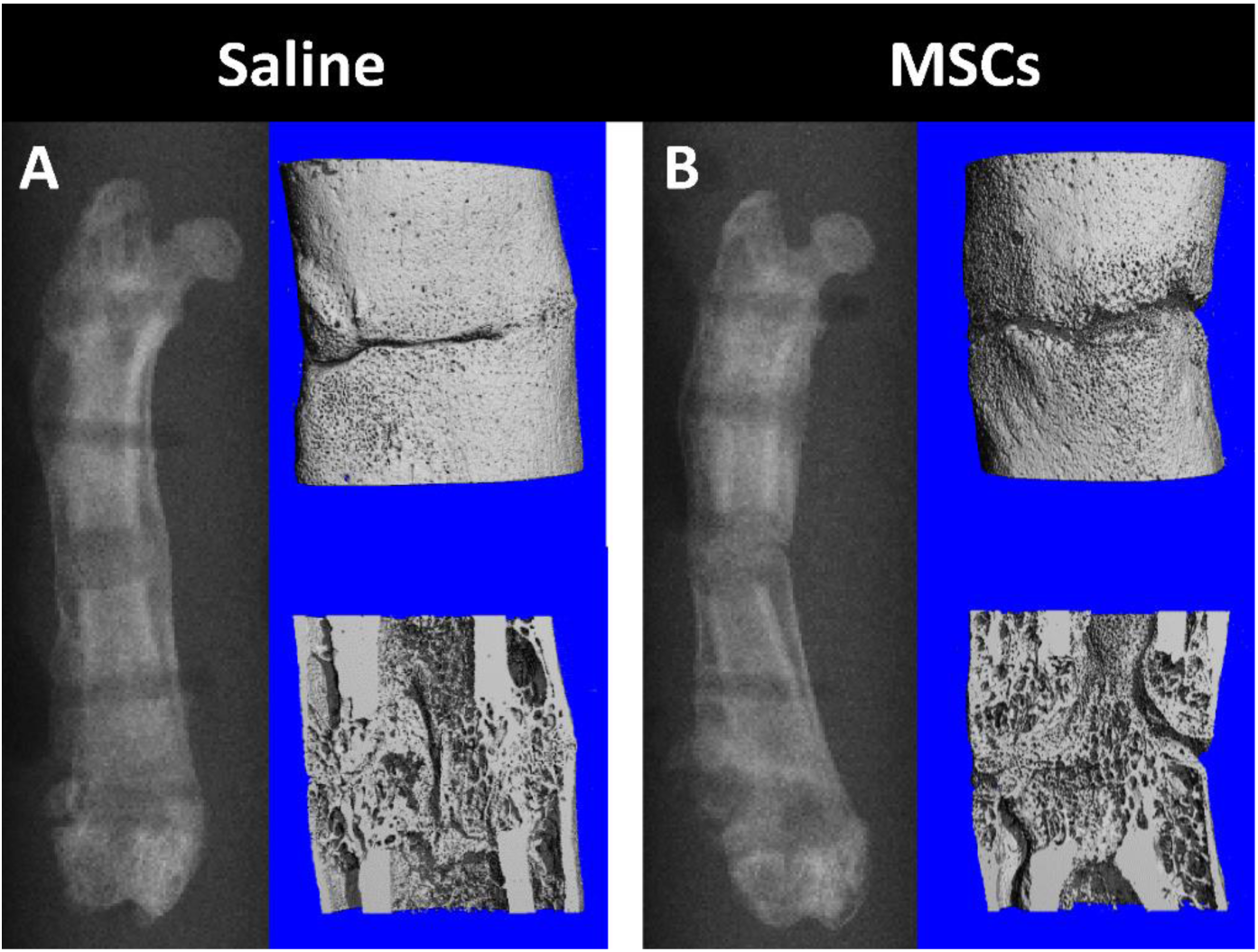
Bone Regeneration. Similar amounts and quality bone was generated in both saline **(A)** and MSC **(B)** treated groups as measured by microCT.

**Fig 4.**
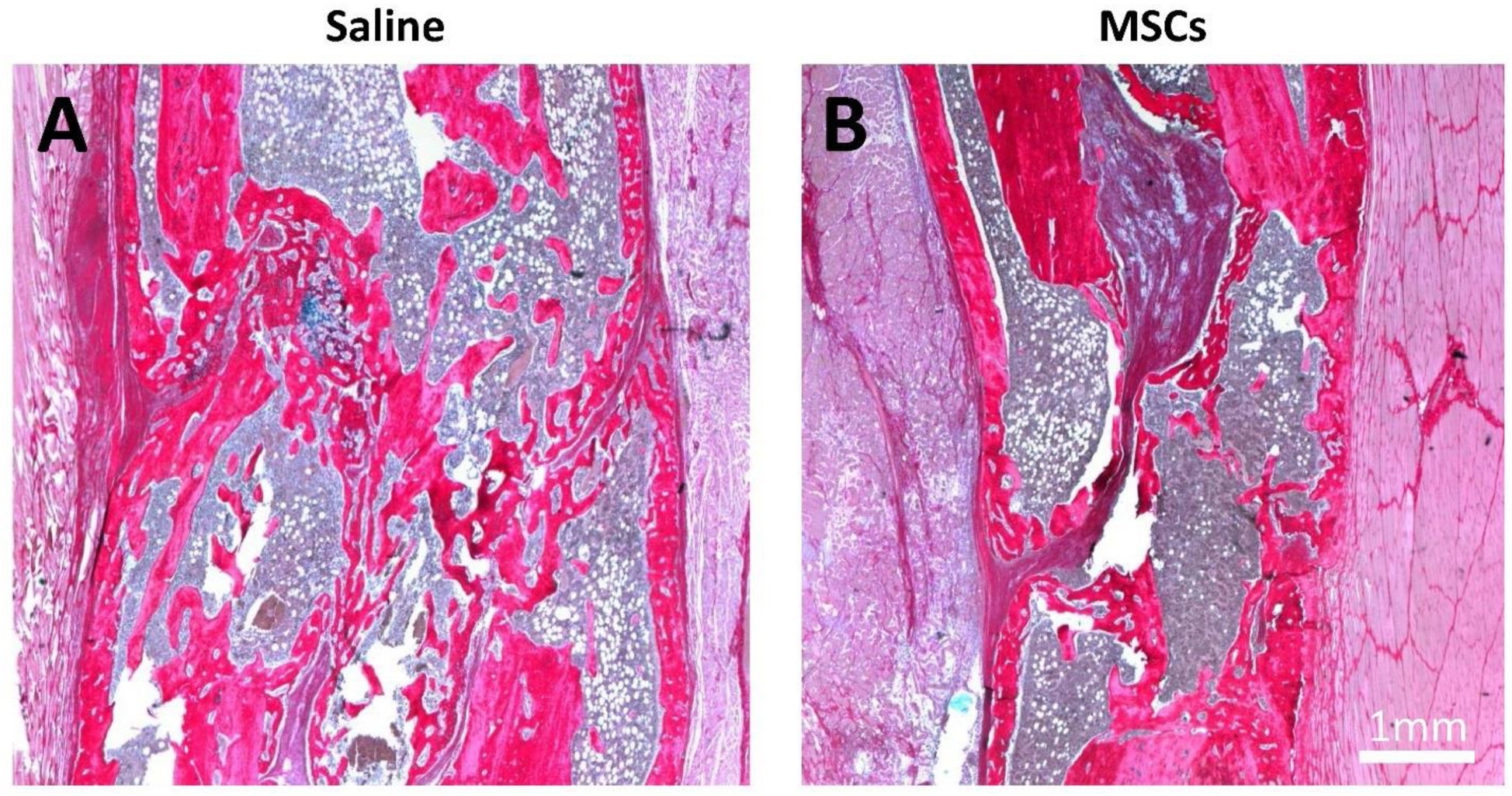
Soft Tissue Outcomes. One section per animal was stained with hemotoxylin, picrosirius red, and alcian blue to differentiate bone (red), fibrous tissue (pink/purple), cartilage (blue), and marrow (grey) in the distraction gap. The relative amounts of each tissue type was similar between saline **(A)** and MSC-treated **(B)** groups.

## Discussion

Mesenchymal stem cells (MSCs) are widely understood to home to sites of injury and have a positive effect.[3,39,42,43] Bone repair is no exception.[28–30,33,35,36,38,39] During complete fracture or drill hole repair, systemically-administered exogenous MSCs will home to the fracture and incorporate into the newly forming tissues.[28–37] Direct implantation of stem cells has been shown to improve the structural and functional outcomes of DO in animal models[12,14,15,17,18,27]. However, there are a number of disadvantages to implementing the same treatment clinically. These could be circumvented if systemic stem cell treatment provides the same results. A previous study has shown that systemically applied MSC’s will home to the distraction site in the mandible similar to other injuries, but is unknown if subsequent bone generation is affected. The purpose of this study was to determine if MSC’s will home to a femoral distraction site and if bone generation is altered. Similar to previous bone repair studies, MSCs did home to the osteotomy site. Although, contrary to our hypothesis, bone regeneration was unaffected.

One possible explanation for the dissimilar outcomes to previous studies is the different models used. Our hypothesis was based on experiments using either complete fractures[30,37] or cortical drill holes.[32–34] Biochemical expression during complete fracture repair and DO has been fairly well characterized[8,44–46] while drill hole has not.[47,48] Early latency, the time of injection for our study, is biochemically very similar to early fracture repair.[8,44–46] A hematoma forms, and inflammatory markers like IL-1 and IL-6 are upregulated to start the repair cascade. A key difference is TNF-α expression which upregulates expression more potently in fracture repair than DO. It has been suggested the TNF-α’s role in bone repair is to recruit additional MSCs in cases of more extensive trauma. Thus, it is possible that although cells were injected at similar numbers at similar times, fewer MSCs homed to or were retained at the distraction site in our study. On the other hand, the active cells and molecules diverge greatly between fracture repair and DO once distraction commences. It is possible that by the time distraction started the exogenous MSCs were committed to cell types incapable of modulating DO-influential molecules.

There are some limitations to our study that need to be considered when interpreting our data. First, only one injection time point was evaluated. Most previous studies have implanted cells into the distraction gap during either the distraction[11,14,20] or consolidation[12,13,15,16,19,23] phases. It is possible that exogenous MSCs do not play a significant role so early in latency, and systemic administration would be effective during later phases. Second, our distraction model heals well without intervention. The animals we used were relatively young and did not have comorbidities like soft tissue damage, diabetes, or smoking that can be present in patients. The final distraction gap was relatively small and did not meet criteria for a critical-sized defect (>5 or 6 mm). The daily distraction rate was in the upper range of what is successful but was safely below the maximum (1mm/day). Since healing is generally achieved without intervention there may not have been room for MSC mediated improvements. It is possible that under more challenging conditions such as an accelerated rate or accompanying soft tissue damage effective MSC contributions could be observed. This was the case in a previous tibia fracture study where healing was inhibited by alcohol administration.[31] Systemic MSC application had little effect on callus volume or strength in non-alcohol administered controls but was able to restore alcohol treated animals to similar morphological and mechanical levels.

Several studies confirm the homing of stem cells in fracture healing and this study sought to take advantage of this homing mechanism to improve bone formation in a DO model. Our results are contrary to previous fracture and direct cell implantation studies. Bone formation was not improved according to our selected outcome metrics. In sum, this does not necessarily mean that systemically applied cells are not beneficial in other ways. More importantly, our results indicate that systemic injection of MSC’s does not harm.

## Acknowledgments

This work was funded by AO North America (Resident Research Grant).

